# Multiple factors influence population sex ratios in the Mojave Desert moss *Syntrichia caninervis*

**DOI:** 10.1101/075861

**Authors:** Jenna T. Baughman, Adam C. Payton, Amber E. Paasch, Kirsten M. Fisher, Stuart F. McDaniel

## Abstract

- *Premise of research:* Natural populations of many mosses appear highly female-biased based on the presence of reproductive structures. This bias could be caused by increased male mortality, lower male growth rate, or a higher threshold for achieving sexual maturity in males. Here we test these hypotheses using samples from two populations of the Mojave Desert moss *Syntrichia caninervis.*
- *Methods:* We used double digest restriction-site associated DNA (RAD) sequencing to identify candidate sex-associated loci in a panel of sex-expressing plants. Next, we used putative sex-associated markers to identify the sex of individuals without sex structures.
- *Key results:* We found an 18:1 phenotypic female: male sex ratio in the higher elevation site (Wrightwood), and no sex expression at the low elevation site (Phelan). In contrast, based on genetic data we found a 2:1 female bias in the Wrightwood site and only females in the Phelan site. The area occupied by male and female genets was indistinguishable.
- *Conclusions:* These data suggest that both differential mortality and sexual dimorphism in thresholds for sex expression likely contribute to population genetic and phenotypic sex ratio biases, and that phenotypic sex expression alone fails to capture the extent of actual sex ratio bias present in natural populations of *S. caninervis.*

## INTRODUCTION

The ratio of males to females in sexually reproducing, dioecious species is generally predicted to be close to 1:1, given the assumption of equal parental investment in offspring of both sexes (Fisher, 1930). Analyses of consistent sex ratio biases therefore have provided important insights into the genetic basis of sex determination or the importance of variation in the roles of males and females in the life cycle (Charnov, 1982; West, 2009). Sex allocation theory predicts that skewed sex ratios can evolve as a result of both competitive and facilitative interactions among relatives (Hamilton, 1967), such that competition for resources among related individuals can bias sex ratios toward the more dispersive sex (Clark, 1978; Gowaty, 1993; Hewison and Gaillard, 1996), while mating among siblings can bias the sex ratio toward females (Herre, 1985; West and Herre, 1998; Reece et al., 2008). The fact that the cytoplasm is generally maternally inherited means that male-killing cyto-nuclear interactions often generate female biases (Schnable and Wise, 1998; Hurst et al., 1999). Other forms of sex-biased mortality also may cause sex ratio biases in specific demographic classes. Thus, generating a clear understanding of consistent sex ratio biases can provide insights into the natural history of a group of organisms, or the factors that govern genetic transmission from one generation to the next. Moreover, because biased sex ratios can profoundly influence the patterns of genetic variation within populations, such biases can have major implications for developing conservation strategies and estimating long term population persistence.

Female biased population sex ratios are a common yet unexplained phenomenon in bryophyte populations (Bisang and Hedenäs, 2005). In dioicous bryophytes, sex is determined at meiosis by a UV chromosomal system (Bachtrog et al., 2011); a spore carrying the female sex-determining locus (U) will form a gametophyte that produces archegonia and ultimately bears the offspring embryo (a sporophyte), while spores with the male sex-determining locus (V) form gametophytes that produce antheridia. Because the diploid sporophyte is produced by the union of a U-bearing egg with a V-bearing sperm, the sporophyte is always heterozygous at the sex-determining chromosome (i.e., UV). As a consequence of meiotic segregation, therefore, the null expectation is a 1:1 sex ratio. Understanding if the persistently female-biased phenotypic sex ratios in moss populations represent a deviation from this genetic expectation is an essential first step toward understanding the ultimate evolutionary causes (e.g., local resource competition, genetic conflict) and consequences of sex ratio bias in this ecologically important group of plants.

Most of the female biases documented in mosses to date have been based on counting the number of sexually mature male and female ramets, or branches, in a population (but see McLetchie et al., 2001; Korpelainen et al., 2008; Bisang et al., 2010, 2014, 2015, Hedenäs et al., 2010, 2016; Bisang and Hedenäs, 2013). A female bias in the production of sexually mature ramets in a natural population could be caused by at least three distinct processes. First, an apparently female biased population sex ratio could simply be a product of faster female growth, as has been found in several species (Shaw and Beer, 1999; Stark, Nichols, et al., 2005; McDaniel et al., 2008). In this case, a population would contain more female ramets than male ramets, and individual female genets (i.e., unique haplotypes) would on average would occupy larger areas than male genets, although the genetic diversity within males and females would be similar (but see Bengtsson and Cronberg, 2009). Second, males might reach sexual maturity less frequently than females. In other words, fewer male haplotypes might produce gametangia than female haplotypes. If this were true, males would constitute a disproportionately large fraction of the sterile plants in a population (termed ‘the shy male hypothesis’; Mishler and Oliver, 1991; Stark et al., 2005). In this case we would expect a genotypic sex ratio that is closer to 1:1 than the sex ratio observed from counting fertile males and females, although again the genetic diversity within males and females would be similar. Finally, a female bias could be caused by elevated male mortality during spore production (McDaniel et al., 2007; Norrell et al., 2014), establishment, or at some later point in the life-cycle. This hypothesis predicts a genotypic female bias at both the ramet and genet level. Regardless of the proximate cause, elevated male mortality would decrease the amount of genetic diversity in males relative to females.

One of the most extreme cases of phenotypic sex ratio variation in mosses is in Mojave Desert populations of *Syntrichia caninervis.* Previous studies based on counts of sexually mature plants indicate that female ramets of *S. caninervis* outnumber males by as much as 7:1 (Paasch et al., 2015) or 14:1 (Bowker et al., 2000; Bisang and Hedenäs, 2005), and that some populations lack sexually mature males entirely (Stark et al., 2001). Mojave Desert *S. caninervis* is extremely desiccation tolerant and spends much of its life in an air-dried state, limiting all biological functions to infrequent post-rainfall periods, primarily in cool winter months (Stark, 1997; Stark et al., 1998). Differences in the timing and duration of this biologically active period, such as precipitation differences along elevation gradients, appear to affect frequency of sex expression (phenotypic sex) in a population. A survey of 890 *S. caninervis* individuals from a 10 hectare elevation gradient (Bowker et al., 2000) found that total percentage of sexually mature individuals increased with elevation. Male sex expression occurred only at the higher elevations, while lower elevation populations contained only a few females with archegonia. In parallel with low levels of sex expression, sexual fertilization and sporophyte production is relatively rare. Established desert *S. caninervis* populations apparently persist through vegetative propagation (Paasch et al., 2015).

Here we expand upon this work using restriction site-associated DNA genome sequencing (ddRADseq) to identify the sex of sterile ramets and study the patterns of genetic variation in males and females in two Mojave Desert populations of *S. caninervis*. We used these data to address four main questions. First, to test whether the phenotypic female bias was caused by higher rates of gametangial production by females, we estimated the phenotypic and genotypic sex ratio in two populations. Second, we estimated genet size to test whether sexual dimorphism in growth rate was sufficient to explain the female biased phenotypic sex ratios in this species. Third, to evaluate whether a genetic sex ratio bias was explained by elevated male mortality, we calculated levels of genetic diversity in male and female haplotypes in both populations. Finally, to evaluate the potential for inbreeding or sex-specific dispersal patterns to generate conditions favorable to local mate competition or local resource competition, we generated several estimates of population differentiation. Collectively these data point to differential sex expression and elevated male mortality as key causes of female-biased sex ratios in Mojave Desert populations of *S. caninervis*.

## MATERIALS AND METHODS

### Sample Collection

In the Mojave Desert, *S. caninervis* grows in a semi-continuous carpet in both shaded and exposed microsites. We identified collections in the field by the characteristic leaf morphology, color, and hair points, although all identifications were confirmed with leaf cross-sections in the laboratory (Mishler, 2007). We collected male, female, and sterile *S. caninervis* samples at two sites from Sheep Creek Wash. The first site (‘Wrightwood’) is located at an elevation of 1800 m in Sheep Creek Wash near Wrightwood, CA (34° 22’ 33.85" N, 117° 36’ 34.59" W), at the west edge of the Mojave Desert and the northern base of the San Gabriel Mountains. The average high and low annual temperature is 16.8 °C and 1.7 °C, with an average annual precipitation of 49.4 cm (2007-2011, Wrightwood Weather Station, NOAA National Climatic Data Center). This site experiences little foot traffic. The second site in the lower portion of Sheep Creek Wash (‘Phelan’) is at an elevation of 1257 m near Phelan, CA (34° 25’ 29.80" N, 117° 36’ 30.91" W), about nine miles northeast of the Wrightwood site. The average high and low annual temperature is 27 °C and 10 °C, while average annual precipitation is 28.2 cm (2005-2009, Phelan, CA, NOAA National Weather Service). This site is also highly disturbed by foot traffic and erosion.

To establish a panel of plants of known sex to use to identify sex-linked molecular markers, we isolated 11 *S. caninervis* female and 10 male ramets from the Wrightwood site in March and April of 2013. In the laboratory under a dissecting microscope, patches were hydrated and screened for presence of current or past antheridia and archegonia.

To estimate the phenotypic sex ratio of the Wrightwood population, in May 2014 we collected in 3-5 cm patches in three parallel 20 m linear north-south (N-S) transects, 10 m apart from one another, collecting one patch every 2 m from a variety of shaded and exposed microhabitats. Collecting at regular intervals along a transect was important for calculating spatial extent of haplotypes and sexes, while collection from different habitat types increased our chances of finding both sexes. In June 2014 we made collections from the Phelan site in a similar manner in two parallel 20 m N-S transects. Additionally, due to the highly irregular distribution of the species in the Phelan site, a third mini-transect was sampled (beginning at about the 23 m mark of transect 2 and extending approximately 2 m), more densely (ca. every 0.3 m) selecting samples in a northwest-southeast (NW-SE) direction from a variety of microhabitats. To estimate the genetic sex ratio and perform diversity analyses, a maximum of three individual sterile ramets were isolated from each patch using a dissecting microscope, resulting in a total of 99 ramets from the Wrightwood site and, due to lower frequency of this species at this site, 42 ramets from the Phelan site, for a total of n = 141 individual ramets.

### DNA Extraction and RADseq library preparation

Total genomic DNA was extracted and isolated from 162 total ramets (21 of known sex, and 141 of unknown sex) using a modified cetyl trimethyl ammonium bromide-beta mercaptoethanol (CTAB) procedure (McDaniel et al., 2007). Prior to extraction, samples were ground dry to a fine powder using a GenoGrinder 2010 bead shaker (SPEX CertiPrep, Metuchen, NJ). DNA quality was evaluated for each sample by electrophoresis and quantified using a Qubit Fluorometer (Invitrogen, Carlsbad, CA, USA), samples’ DNAs were normalized prior to library preparation.

Illumina libraries were prepared for sequencing using a modified version of the Peterson et al. (2012) protocol using the endonucleases EcoRI and MseI (New England Biolabs, Ipswich, MA). Following double enzyme digestion, unique barcoded adaptors were ligated to the resulting EcoRI cut sites and a non-barcoded universal adaptor to the MseI cut site. Variable length barcodes of 8, 9, and 10 base pairs (bp) were used with each barcode, differing by at least 4 bp. Illumina flow cell binding and sequencing primer sites were added to the adaptor ligated fragments throughout 10 cycles of PCR using NEB Q5 PCR master mix (New England Biolabs, Ipswich, MA). Success of library construction was evaluated through agarose gel electrophoresis, after which 5 µL of each sample’s final library was pooled into a single tube for the 141 sterile samples and a separate tube for the 21 samples of known sex. Size selection and sequencing were performed at the University of Florida’s Interdisciplinary Center for Biotechnology Research. Pooled libraries were fractioned using Pippin ELF precision electrophoresis (Sage Science, Beverly, MA) with the resulting 250-500 bp fraction being used for Illumina sequencing. The sterile ramet sample library was sequenced using an Illumina NextSeq 500 at mid throughput, producing 150 bp single-end reads. The library consisting of the 21 known-sex ramets was pooled with libraries constructed in an identical manner for a non-related study and sequenced on one lane of a HiSeq 3000 producing 100 bp single end reads.

### Data analysis

Raw ddRAD sequence reads were assessed for quality with fastQC (Gordon and Hannon, 2010). All manipulation of raw sequence reads was performed using tools from the FASTX-Toolkit (Gordon and Hannon, 2010) implemented in Galaxy on the University of Florida’s Research Computing Cluster. To eliminate low quality bases at the 3’ end of reads, all sequences were trimmed to 100 bp. Reads were filtered for a minimum Phred quality score of 20 on at least 90% of the read. All reads were then demultiplexed followed by the removal of the 5’ barcode and EcoRI enzyme cut site.

RAD loci were assembled de novo using STACKS, version 1.24, (Catchen et al., 2011, 2013). Pertinent parameters for the STACKS pipeline were as follows: a minimum of 2 reads required for each allele within an individual (-m option ustacks), a maximum allowable distance of 2 nucleotides between alleles within a locus (-M option ustacks), a maximum of 2 mismatches allowed when aligning secondary reads to primary stacks (-N option ustacks), and a maximum nucleotide distance of 2 allowed between loci of different individuals (-n option cstacks). A subset of 30 samples, 15 from each site, was used to create the master catalog of loci. Using the STACKS ‘populations’ program, loci containing any SNP categorized as heterozygous for an individual were identified. Since all individuals sampled were haploid gametophytes, all loci were expected to be homozygous; therefore, those loci identified as heterozygous were discarded as they likely represented paralogous or over-merged loci. Approximately 23% of loci had at least one individual with a heterozygous SNP call. Although removing only the heterozygous allele may be sufficient for removing errors, to be conservative we remove the entire locus from the dataset. Sequence data were further filtered to retain only bi-allelic loci with 1 or fewer SNPs, found in a minimum of 74% of samples, with a minor allele frequency lower limit of 0.1, resulting in 2,234 loci used for subsequent genetic analyses.

Twenty-one *S. caninervis* ramets of known sex (11 females expressing archegonia and 10 males expressing antheridia) were used to identify potential sex-associated loci (fragments) that would allow us to infer sex of sterile ramets. Any 80 bp locus that was present in most individuals of one sex and absent in all individuals of the opposite sex was considered a potential sex-associated locus. Using these criteria, we identified over 5,000 candidate sex-associated loci. Although one sex-linked locus would suffice for our purposes, we selected the 100 male-associated and 100 female-associated loci with the best coverage across all 21 individuals for further analyses. This is because a single autosomal locus may show a spurious association with sex in a sample of that size, due to chance or because it has a sex-biased allele frequency; the region of suppressed recombination on the UV sex chromosomes, however, makes it likely that many sex-linked markers will segregate in complete linkage disequilibrium, providing a signature of true sex linkage. We used a custom Perl script to search for the candidate sex-linked sequences in the alignments from the final 131 sterile ramets used in our analyses (see Results). We then used the loci that were in complete linkage with one another in the 131 sterile ramet samples to determine the sex of each haplotype and tested for deviations from a 1:1 sex ratio with a Chi-square test and a 2-tailed p-value.

In order to determine the number of unique haplotypes among the 131 sterile ramets, clonal assignments were made using genetic distance under the infinite allele model with a pairwise distance threshold of 0 between individuals. Higher mismatch thresholds, up to 8, were evaluated but all resulted in the same clonal assignments. Figure 1 shows the clonal decay distribution of number of unique genets identified at different genetic distance threshold levels. Furthermore, all ramets that were identified as clones of the same genet were also genetically assigned to the same sex. Clonal assignments were used to fill in missing data, when possible, by finding a consensus sequence among ramets per haplotype using GENEIOUS version 8.1.4 (Kearse et al., 2012). To test whether the female biased sex ratio was generated by greater female growth rates, we compared the mean number of sites occupied by female and male haplotypes, as well as the mean number of ramets per male and female haplotypes, testing for significance using a one-tailed t-test.

**Figure 1.**
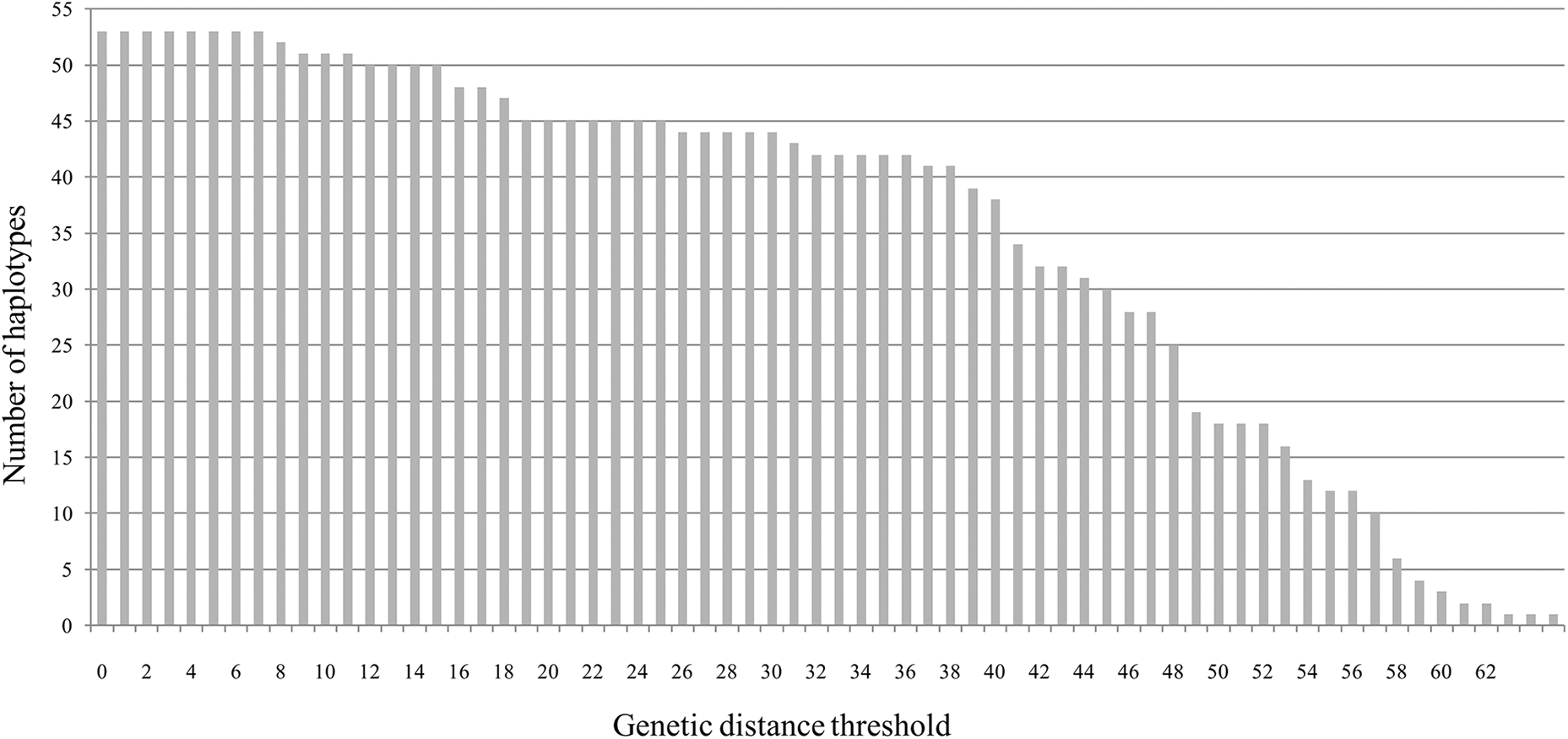
Clonal lineage decay at different genetic distance thresholds. This figure shows the number of genets that would be identified at each genetic distance parameter setting.

To compare levels of genetic variation between the two study populations, and between males and females in the Wrightwood population, we calculated five measures of clonal diversity using the software GENODIVE version 2.0b23 (Meirmans and Tienderen, 2004). Clonal diversity (P_d_) is a measure of unique genets relative to (divided by) the number of ramets sampled (Ellstrand and Roose, 1987). The effective number of genotypes (N_eff_) is an index that accounts for frequencies of each genet and describes the number of haplotypes that have equal frequencies while minimizing low frequency haplotypes. This index is analogous to effective number of alleles but instead counts whole haplotypes. Clonal evenness (effective number of genotypes divided by number of genotypes) indicates how evenly the haplotypes are divided over the population and would be equal to 1.00 if all haplotypes were represented equally. We also calculated Simpson’s diversity index, corrected for sample size, and the corrected Shannon’s index (Chao and Shen, 2003). The latter is a measure of clonal variation that accounts for singletons (genets or haplotypes sampled just once) in the population for sample sizes greater than approximately 50 sampling units (Arnaud-Haond et al., 2007). We used bootstrap tests with 1,000 permutations and subsampling to match population sizes to test for differences in the latter three measures of clonal diversity.

To evaluate the degree to which differences between the two populations could be due to allele frequency differences, we first calculated the population differentiation statistic F_ST_ among the 131 sterile ramets from the Phelan and Wrightwood sites using

GENODIVE. Next, we estimated the population structure using the fastSTRUCTURE (Raj et al., 2014) inference algorithm with a simple logistic prior and K=1 through K=4. The dataset used for this contained 2,234 SNPs from 131 sterile ramets from the Phelan and Wrightwood sites, where missing data were filled in with data from identical clonal haplotypes, when possible. Membership coefficients for K=2 were plotted using DISTRUCT version 2.2 from the fastSTRUCTURE software package.

For another means to visualize the patterns of genetic distance among genets in the two populations, we constructed a midpoint-rooted neighbor joining genetic distance tree using CLEARCUT. The dataset used for this contained 2,234 SNPs from 51 genets from the Phelan and Wrightwood sites, where missing data were again filled in with identical haplotypes, when possible. Two genets with greater than 80% missing data were excluded. Because the RAD loci are not completely linked, this tree does not represent the genealogical relationships among these individuals but rather genetic similarity.

## RESULTS

### Phenotypic sex ratios

Of the 49 patches collected in the Wrightwood site, 31 contained no sex structures, 17 contained ramets expressing archegonia and were classified as female, one contained ramets of both sexes, and no patches contained ramets with only male gametangia. The resulting phenotypic sex ratio of 18 F:1 M differs significantly from 1:1 meiotic expectations (Chi-squared test, two-tailed p-value < 0.0001).

### Sequencing statistics

Approximately 150 million total raw sequencing reads were generated for the 141 ramet samples sequenced resulting in roughly 1.06 million reads per barcode. About 40% of the reads were discarded due to low quality, resulting in 88 million reads that passed initial quality filters. Ten ramets with fewer than 4.5 thousand reads were discarded, leaving 98 sterile ramets in the Wrightwood site, 33 sterile ramets in the Phelan site, and 21 ramets of known sex from the Wrightwood site, ranging from 51 thousand to 2.6 million reads per ramet. The data matrix of 2,234 SNPs from 131 sterile ramet samples was 80% complete when using original reads but increased to 94% complete when missing data were filled in with data from identical clonal haplotypes. The data matrix of concatenated loci used for the RAxML analysis was 83.3% complete with a mean depth of about 5.6 reads per locus.

### Identifying sex linked loci and genetic sex ratio

We used the 20 individuals of known sex to identify 100 candidate male and 100 candidate female diagnostic markers. We chose this approach because our sample size was necessarily small, meaning that any single locus could show a spurious or population specific linkage to sex. Indeed, approximately 25% of the loci from our trial set (52 loci between males and females) were absent from our individuals of unknown sex, and 10% of the loci were apparently misclassified as sex-linked because they failed to show complete linkage disequilibrium with the other candidate loci. It is possible that the missing loci were lost due to subtle differences in the library size selection procedure, or due to stochastic sampling during the sequencing process. The remaining 65% of the candidate sex-linked loci were in complete linkage with one another, as expected for a UV sex chromosome system. The 63 putatively male-linked loci had an average read depth of 6.3 and on average 33.5 male-associated loci were found within each newly identified genotypic male ramet. The 65 putatively female-linked loci had a mean read depth of 5.7 and an average of 25.3 female-associated loci were detected in each newly identified genotypic female ramet. Thus, although some of these loci may later be shown to be only sex-correlated, or limited to these populations, and not truly sex-linked, these data give us high confidence that collectively these loci are diagnostic markers for sex in this sample of individuals.

Using these putative sex-linked markers, we found 65 female ramets and 33 male ramets in samples from the Wrightwood site, equating to an approximate 2:1 female-biased genetic sex ratio. This represents a significant deviation from both the 1:1 expectation (Chi-squared test, two-tailed p-value = 0.0012) and from the observed 18:1 phenotypic sex expression based on counts of sexually mature ramets (Chi-squared test, two-tailed p-value < 0.0001). All 33 of the ramets from the Phelan site were female.

Including all of the 2,234 biallelic SNPs, we found 53 distinct genetic individuals, or haplotypes, among the 131 sterile ramets. The Wrightwood site contained 45 unique haplotypes and the Phelan site had eight haplotypes. A total of 53 haplotypes were distinguishable at genetic distance thresholds of 0 through 7, after which the number of haplotypes began to drop incrementally without distinct breaks, indicating that the number of genotypes is unlikely to be inflated as a result of genotyping errors (Figure 1). The plants in the Wrightwood site were assignable to 31 female haplotypes and 14 male haplotypes, corresponding to an approximately 2:1 female-biased sex ratio.

### Distribution of genotypic variation

To test whether males and females exhibited different patterns of vegetative expansion, we counted the number of sites that each haplotype occupied. Most haplotypes in the Wrightwood site were restricted to a single 3-5 cm patch. Of the 45 haplotypes in the Wrightwood site, three were found within adjacent patches (approximately 2 m apart) and only one haplotype spanned three patches across about 4 m (Figure 2). Female and male haplotypes did not occupy significantly different numbers of patches (mean female patches = 1.097 ± 0.301, mean male patches = 1.143 ± 0.535, one-tailed t-test p-value = 0.3565) nor did they contain significantly different numbers of ramets per haplotype (mean female ramets = 2.097 ± 1.044, mean male ramets = 2.357 ± 1.151, one-tailed t-test p-value = 0.2286).

**Figure 2.**
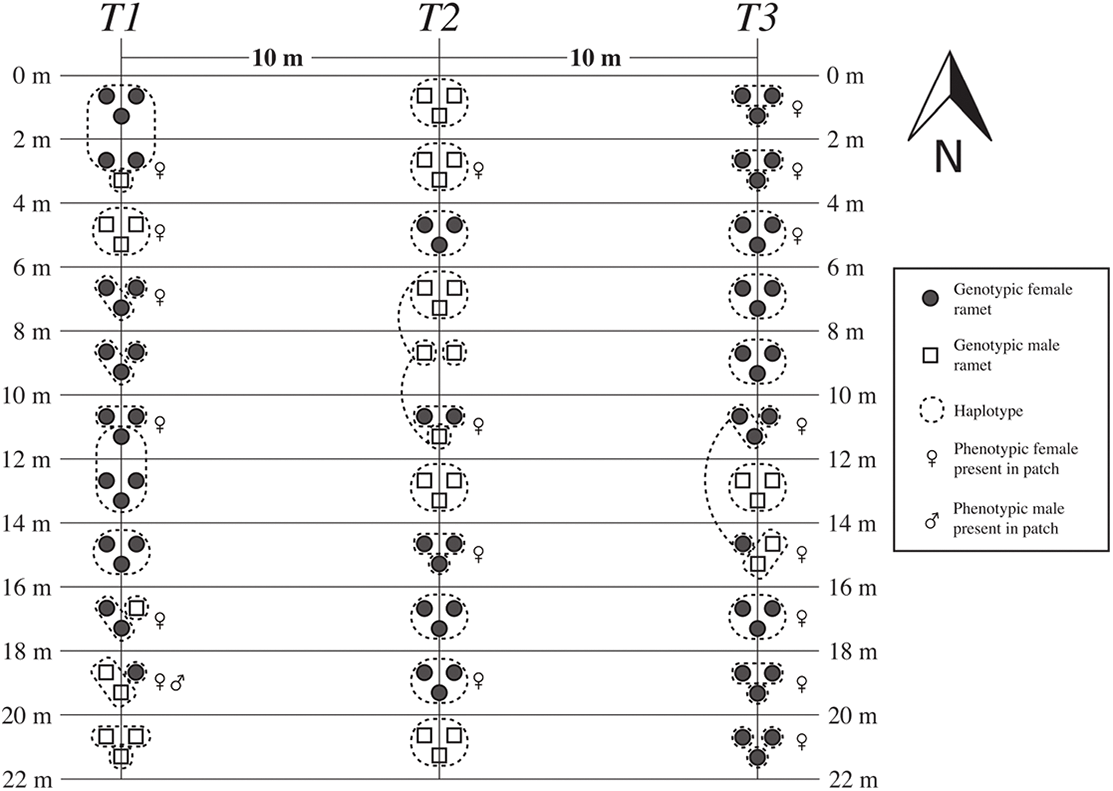
Spatial distribution of genets. Wrightwood contained 53 unique genets in 98 ramets from 33 patches along three transects (T1, T2, and T3). Genotypic females are represented as circles and genotypic males as squares. Genets are outlined in dashed lines. In cases that genets are in different patches, a dashed line connects different ramets of the same MLG. Presence of phenotypic sex in each patch is indicated with ♀ for phenotypic females and ♂ for phenotypic males.

To test whether the sexes harbored different amounts of genetic diversity, we calculated several indices. The Simpson’s diversity and evenness values were not significantly different between the two sexes in Wrightwood, but the corrected Shannon Index was significantly higher for females than for males (Table 1, p-value = 0.001). Both the Simpson’s diversity and the corrected Shannon indices were significantly higher in Wrightwood than in Phelan (p-value = 0.001). Evenness was also slightly greater in the Wrightwood site, but not significantly so (Table 2).

**Table 1.** Clonal diversity indices for the sexes in the Wrightwood site.

| <u>Sex</u> | <u>N</u> | <u>G</u> | <u>P<sub>d</sub></u> | <u>N<sub>eff</sub></u> | <u>Simpson's</u> | <u>Evenness</u> | <u>Shannon (corrected)</u> |
| --- | --- | --- | --- | --- | --- | --- | --- |
| Female | 65 | 31 | 0.477 | 25.000 | 0.975 | 0.806 | 1.596* |
| Male | 33 | 14 | 0.424 | 11.463 | 0.941 | 0.819 | 1.208* |

Note: \*Statistical difference between the sexes in Wrightwood, p-value = 0.001, based on 1,000 permutations with subsampling to match population sizes. N = number of ramets sampled, G = number of genets, P<sub>d</sub> = clonal diversity, N<sub>eff</sub> = effective number of genotypes.

**Table 2.** Clonal diversity indices for Wrightwood and Phelan sites.

| <u>Site</u> | <u>N</u> | <u>G</u> | <u>P<sub>d</sub></u> | <u>N<sub>eff</sub></u> | <u>Simpson's</u> | <u>Evenness</u> | <u>Shannon (corrected)</u> |
| --- | --- | --- | --- | --- | --- | --- | --- |
| Wrightwood | 98 | 45 | 0.459 | 36.379 | 0.983* | 0.808 | 1.753* |
| Phelan | 33 | 8 | 0.242 | 6.368 | 0.869* | 0.796 | 0.874* |

Note: \*Statistical difference between Wrightwood and Phelan, p-value = 0.001, based on 1,000 permutations with subsampling to match population sizes. N = number of ramets sampled, G = number of genets, P<sub>d</sub> = clonal diversity, N<sub>eff</sub> = effective number of genotypes.

### Population differentiation and structure

To evaluate the potential for sex biased dispersal or inbreeding to drive sex ratio bias, we examined several measures of population structure. The F_ST_ between the Phelan and Wrightwood sites was 0.102 when all ramets were included. When estimated with just unique haplotypes, however, F_ST_ dropped to 0.028, indicating that the patterns of haplotype expansion differ in the two populations. The neighbor joining tree recovered haplotypes collected from Phelan nested within Wrightwood haplotypes (Figure 3). Population structure analysis in fastSTRUCTURE estimated a minimum of one model component to explain structure in the data. However, this estimate was not supported by the corresponding marginal likelihoods, which increased with the K parameter with no apparent maximum.

Membership coefficients for K =2, visualized on Figure 4 show some support for structure among the two sites, consistent with inferences based on F_ST_.

**Figure 3.**
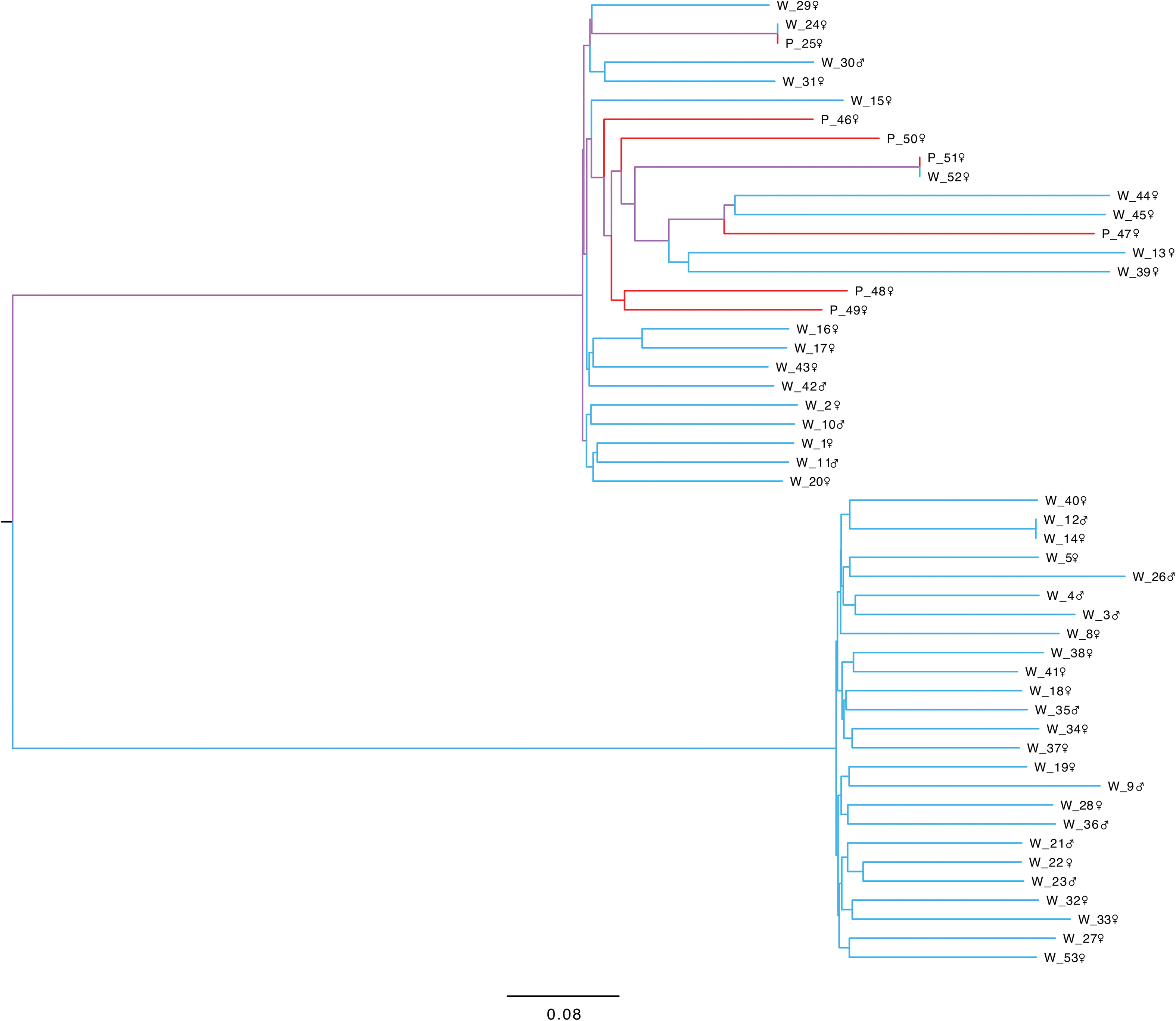
CLEARCUT neighbor joining genetic distance tree of ramets from the Wrightwood and Phelan sites. Branches that join samples from the Wrightwood site (‘W’) site are blue and those that join the Phelan site (‘P’) are red. Branches that support mixed-population groups are purple. Symbols indicate inferred sex.

**Figure 4.**
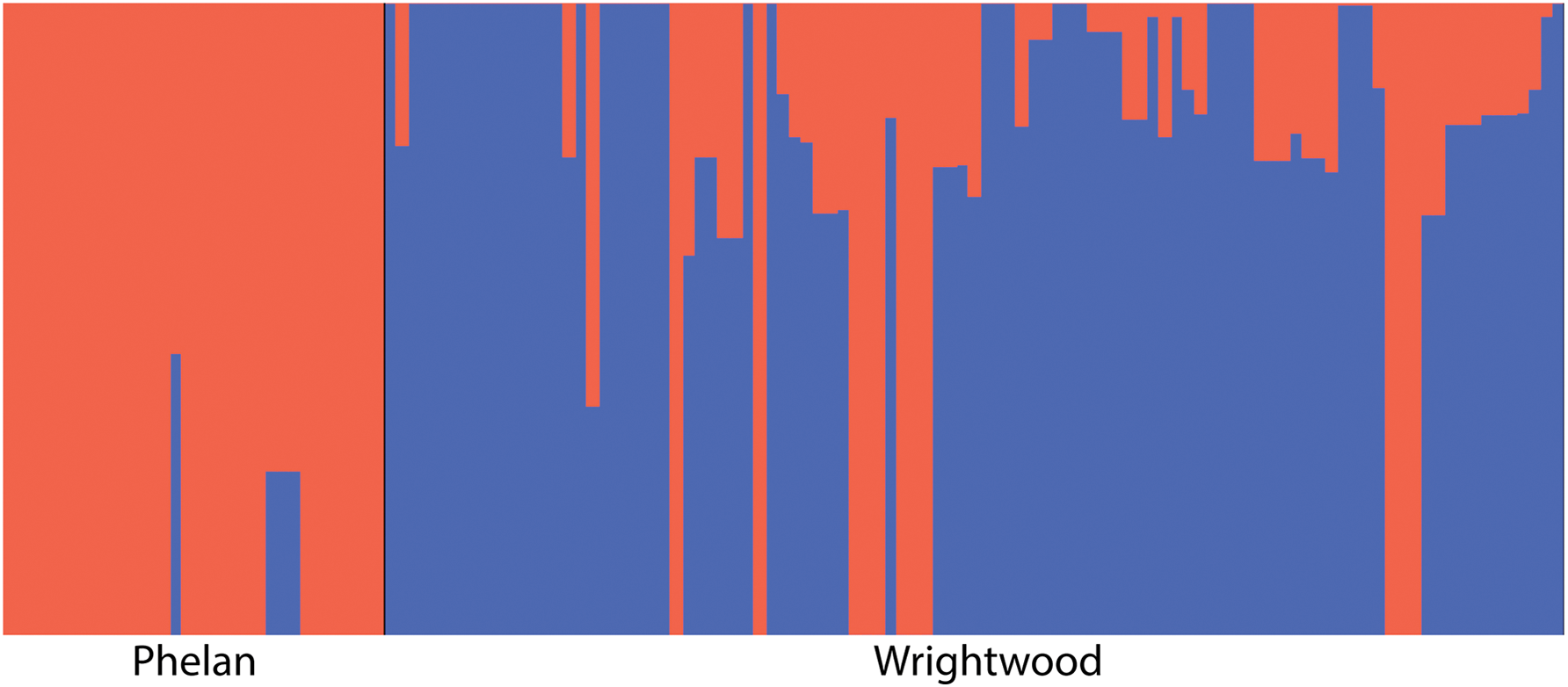
fastSTRUCTURE membership coefficients for K = 2. Vertical bars represent individual ramets while colors represent genetic clusters detected (K = 2). Bars on the leftrepresent ramets from the Phelan site (n = 33) and bars on the right represent ramets from the Wrightwood site (n = 98). Membership coefficients are plotted for each ramet and are colored corresponding to the proportion of each ramet MLG that most closely aligns with each of the two clusters. The dataset used for this analysis contained 2,234 SNPs from 131 sterile ramets from Wrightwood and Phelan where missing data were filled in with clones, when possible.

## DISCUSSION

*S. caninervis* has long been a model for investigations regarding the evolution and ecology of sex ratio variation in mosses (Stark et al., 1998, 2001; Stark, McLetchie, et al., 2005; Stark, Nichols, et al., 2005; Stark and McLetchie, 2006; Paasch et al., 2015).

However, our understanding of sex ratio variation in this species, as well as other bryophytes, has been limited by the large number of sterile plants in most bryophyte populations. In principle, this limitation is overcome by the use of sex-linked molecular markers, which provide a simple means of determining the sex of sterile plants. Identifying reliable sex-linked markers is not trivial, though, requiring screening large numbers of polymorphic loci in a large pedigree or mapping population. In addition, sex linkage ideally should be confirmed using a physical map, or molecular evolutionary analyses, for example to test for complete linkage disequilibrium (LD) among sex linked loci, as predicted based on the observation that recombination does not occur on UV sex chromosomes (Bachtrog et al., 2011). To date, within mosses this has only been accomplished in the model system *Ceratodon purpureus* (McDaniel et al., 2007, 2013). A less rigorous approach involves testing for an association between a molecular marker and sex in a large panel of unrelated individuals (as has been done in three *Drepanocladus* species; (Bisang et al., 2010, 2015; Bisang and Hedenäs, 2013; Hedenäs et al., 2016). Here, we have used the latter approach, and screened several thousand restriction-site associated sequenced (RADseq) loci in a small number of individuals of known sex. Although the number of individuals was small, the large number of loci gave us additional confidence because we could retain only those that exhibited complete LD among all loci in a larger sample. Beyond the large numbers of polymorphisms, RADseq loci have the advantage of being defined by a unique DNA sequence, which both allows us to be confident in the homology of our loci (unlike gel band-length approaches), and ultimately provides a means of identifying sex-linked genes in published transcriptomes (Gao et al., 2014; Wickett et al., 2014).

We use these putative sex-linked markers to show that the genotypic sex ratios in two Mojave Desert populations populations of *S. caninervis* are female biased (2 F:1 M in the Wrightwood site, and entirely female in the Phelan site). Importantly, however, the phenotypic sex ratios in this sample were far more biased (18 F:1 M in Wrightwood). While it is certainly possible that we have over-estimated the long-term female sex expression bias, due to drought in the study area in the years prior to collection (2013-2014), collections from seasons with more typical winter weather patterns reported phenotypic sex ratios of 7 F:1 M (Paasch et al., 2015), approximately three times more female biased than the genotypic sex ratio we found. These data indicate that males constitute a disproportionately large fraction of the sterile plants, providing unequivocal support for the shy male hypothesis in these Mojave Desert populations of *S. caninervis*.

The greater frequency of female ramets, however, indicates that factors beyond thresholds for sex expression must also contribute to the population sex ratio variation in *S. caninervis*. Experimental manipulations show that females regenerate more readily from plant fragments than males do under both cool conditions and desiccation stress (Stark et al., 2004; Stark, Nichols, et al., 2005; Stark and McLetchie, 2006), and potentially may accumulate more biomass. However, the fact that we observed no difference in number of sites occupied between female and male haplotypes indicates that the genotypic female bias in these populations is unlikely to be due to faster female growth.

We do, however, find some evidence that elevated male mortality contributes to the overall female biased sex ratio. The male-mortality hypothesis uniquely predicts a lower male haplotype diversity, relative to females, as we found (corrected Shannon index, 1.596 in females, 1.208 in males, p = 0.001). When comparing the two populations, the Phelan site is less clonally diverse (Simpson’s and corrected Shannon indices, p = 0.001) and has a greater genotypic female bias than the Wrightwood site, also suggesting greater mortality. Importantly, the relatively low F_ST_ between these two populations indicates that most surveyed polymorphisms are shared between Phelan and Wrightwood, suggesting that sex ratio differences between them are unlikely to result from long isolation.

With our current data it is not possible to isolate when in the life cycle male and female demography differ, nor whether the two sexes exhibit different microhabitat preferences. However, the observed pattern of lower male diversity (i.e., elevated male mortality) suggests that the interaction between male physiology and extrinsic environmental factors are more likely to govern population sex ratio than exclusively intrinsic factors like sex ratio distorters (McDaniel et al., 2007; Norrell et al., 2014). Indeed, the available evidence suggests that *S. caninervis* female biased phenotypic sex ratios in Mojave Desert populations correlate with precipitation and temperature (Bowker et al., 2000). One potential cause of the rarity of males observed in *S. caninervis* is the difference in timing of resource allocation to reproduction between the sexes. Although sexual reproduction is costly for both sexes, in the moss *Hylocomium splendens* males make a higher initial investment in the production of antheridia, while fertilized females only allocate energy to reproduction after fertilization through nurturing a sporophyte (Rydgren et al., 2010). Although we observe sporophytes only rarely in *S. caninervis*, the relatively high diversity evident in our sample, along with the limited population structure and weak structure in the population genealogy all indicate that sexual reproduction is or has been present in these populations. Thus, males may experience a trade-off between sexual selection, to produce more reproductive tissue, and natural selection, which may favor investment in the maintenance of vegetative tissues, as has been seen in the moss *C. purpureus* (McDaniel, 2005).

Several bryophyte demographic models predict the eventual local extinction of males, based on vegetative growth patterns similar to those found in *S. caninervis* (McLetchie et al., 2001; Crowley et al., 2005; Rydgren et al., 2010), although these models generally assume that migration is negligible. The lack of population structure between our two study sites indicates that population sex ratios may be influenced by migration of spores from other populations in addition to local population dynamics. The mixing of genotypes in the distance tree suggests that both sites in Sheep Creek Wash draw spores from the same metapopulation. Some populations, like the Wrightwood site, both recruit and produce spores that contribute to subsequent generations, while other sites, like Phelan, permit growth but no sexual reproduction and become genotype sinks. Our sampling is insufficient to quantitatively evaluate the population structure of Mojave Desert populations of *S. caninervis*, but these inferences are consistent with studies in other mosses that report some population structure at the patch scale (Hutsemékers et al., 2010; Leonardía et al., 2013; Rosengren et al., 2016) but limited structure at regional or larger spatial scales (McDaniel and Shaw, 2005; Vanderpoorten et al., 2008; Korpelainen et al., 2012; Shaw et al., 2014; Désamoré et al., 2016).

### Conclusions

This study demonstrates that the highly female-biased sex ratios observed in Mojave Desert *S. caninervis* are congruent with both the shy male hypothesis (Mishler and Oliver, 1991; Stark, McLetchie, et al., 2005) and increased male mortality (Stark et al., 2000). These results suggest the importance of interactions between environmental conditions and demographic factors for shaping sex ratios in this species, and may have important consequences for the persistence of local populations in the presence of long-term shifts in climate. Importantly, both mechanisms are grounded in the disproportionate pre-zygotic energetic allocation to sexual reproduction by males relative to females (Mishler and Oliver, 1991; Stark et al., 2000; Stark, McLetchie, et al., 2005), in essence representing a trade-off between natural and sexual selection. Our data suggest that local mate competition and local resource competition are less likely to be major drivers of sex ratio variation in Mojave Desert populations of *S. caninervis*, but we should caution that studies of inbreeding and sex-biased dispersal are needed in bryophytes. We suspect that the wider use of genomic approaches like those we have used here is likely to uncover cases of sex ratio variation where other processes may predominate (Cronberg, 2003; Bisang and Hedenäs, 2005; Stark et al., 2010; Horsley et al., 2011; Bisang et al., 2014; Norrell et al., 2014). A future challenge is to determine the contributions of migration and environmental factors like water availability, through its effects on determining sex expression and mortality, on the trajectory of population sex ratio change through time.

## Acknowledgements

This work was supported by the California State University Department of Biological Sciences Faculty-Student Achievement Fund to KMF, a Research Exchange Grant to JTB from the NSF EDEN RCN (IOS 0955517), the NSF Louis Stokes Alliances for Minority Participation (HRD-1363399) to AEP, the NIH-NIGMS Minority Biomedical Research Support-Research Initiative for Scientific Enhancement program (GM61331) to JTB, and an NSF grant (DEB 1541005) to SFM.

## APPENDIX 1

Specimen vouchers were deposited in the California State University, Los Angeles Herbarium (CSLA).

